# Luteolin transforms the BMDM polarity to regulate the expression of inflammatory factors

**DOI:** 10.1101/2020.06.30.181503

**Authors:** Shuxia Wang, Shuhang Xu, Meng Cao, Jing Zhou, Xiaodong Mao, Xiaoming Yao, Chao Liu

## Abstract

Macrophage are indispensable regulator cells in inflammatory response. Macrophage polarization and its secreted inflammatory factors have affinity with the outcomes of inflammation. Luteolin, a flavonoid abundant in plants has anti-inflammatory activity, but whether luteolin can manipulate M1/M2 polarization of BMDM to suppress inflammation is still veiled. The purpose of this study was to observe the effects of luterolin on the polarity of BMDM derived from C57BL/6 mice and the expression of inflammatory factors, to explore the mechanism of luteolin regulating the BMDM polarity. M1-polarized BMDM were induced by LPS+IFN-γ, M2-polarization were stimulated with IL-4. BMDM morphology was observed by laser confocal microscopy; levels of BMDM differentiation and CD11c or CD206 on membrane surface were assessed by FCM; mRNA and protein of M1/M2-type inflammatory factors were performed by qPCR and ELISA, respectively; the expression of p-STAT1 and p-STAT6 protein pathways was detected by Western-blotting. The isolated mouse bone marrow cells were successfully differentiated into BMDM, LPS+IFN-γ induced BMDM M1-phenotype polarization, and IL-4 induced its M2-phenotype polarization. After M1-polarized BMDM treated with luteolin, M1-type pro-inflammatory factors including IL-6, TNF-α□iNOS, CD86 were down-regulated while M2-type anti-inflammatory factors including IL-10, Arg1, CD206 were up-regulated; the expression of M1-type surface marker CD11c decreased, nevertheless, M2-type marker CD206 increased; levels of inflammatory signaling protein p-STAT1 and p-STAT6 were attenuated and enhanced respectively. Our study suggests luteolin may transform BMDM polarity through p-STAT1/6 to regulate the expression of inflammatory mediators, thereby inhibiting inflammation. Naturally occurring luteolin hold promise as an anti-inflammatory and immunomodulatory agent.

---

Inflammation is the immune system’s response to invading pathogens, but aberrant inflammation responses leads to a “cytokine storm” that makes patients sicker (1). Mounting evidences found that continuous and/or repeated inflammatory stimuli could also induce tumors (2). Therefore, the inflammatory response is a double-edged sword. If immune cells and pro-inflammatory cytokines are overproduced, cytokine cascades occur, called “cytokine storm” or termed as “inflammatory storm”, leading to sepsis, acute respiratory distress syndrome (ARDS) and even multiple organ failure (MOF) (3). As well all know, pathogenic agents such as viral or bacterial infections incur the pathological process of sepsis which is characterized by an overwhelming generation of pro-inflammatory cytokines. Recently, global pandemic of coronavirus disease 2019 (COVID-19) suffering from severe acute respiratory syndrome coronavirus 2 (SARS-CoV-2) is also associated with macrophage hyperpolarization elicits “cytokine storms” and viral sepsis (4). Thus, the immunomodulatory therapy of inflammation is crucial for maintaining homeostasis (5-6).

Macrophages have been identified as critical effector cells in inflammatory/immune response and can be activated by pathogenic agents or inflammatory mediators to secrete various inflammatory factors. Meanwhile, heterogeneity and plasticity are hallmarks of macrophages, that is, M1-polarized (pro-inflammatory) macrophages and M2-polarized (anti-inflammatory) macrophages. Pathogens infection can polarize macrophages to M1-phenotype, produce high level of pro-inflammatory cytokines such as IL-6 and TNF-α, or effector molecules iNOS and surface markers CD11c or CD86, exert a pro-inflammatory effect and defense against pathogens. Conversely, IL-4 or TGF-β induced M2-phenotype macrophages mainly express anti-inflammatory cytokine IL-10, effector molecule Arginase (Arg) 1 and surface marker CD206, contribute to hinder inflammation (7). Normally, the M1/M2 polarization of macrophages maintains a dynamic equilibrium. When virulent bacteria or viral infections or overmuch inflammatory molecules irritation, this balance is disrupted, excessive M1 polarization of macrophages will generate redundant inflammatory factors, causing systemic inflammatory response syndrome (SIRS, namely sepsis) and MOF (8). Therefore, it is particularly vital to skew the macrophage polarization and avoid excessive M1-polarization, thus reduce the inflammatory response and promote tissue remodeling.

At present, most anti-inflammatory agents are glucocorticoids, antibiotics or antivirals. Hormones not only produce immunosuppression, but also induce secondary infections and prolong the disease course or other side effects. Antibiotics or antiviral drugs only kill the pathogen, and that antibiotics lyse the bacteria while killing the bacteria, releasing more toxins to induce “cytokine storms”, further exacerbating the inflammatory response and promoting the promoting factor to the development of sepsis. In this regard, natural anti-inflammatory immune drugs extracted from plants have multi-effect regulation and less toxic, and especially have obvious benefits in suppressing inflammation. Therefore, it is urgent to seek natural compounds with high efficiency and low toxicity as anti-inflammatory immune agents.

Luteolin (Lut) is a flavonoid, mainly exists in fruits, vegetables and Chinese herbs, which has anti-hyperlipidemia (9), antitumor (10), anti-inflammation and immunoregulation (11). Studies showed that luteolin also has antiviral effects against Influenza A virus or dengue virus by interfering with coat protein (12-13). Recently research indicated that active ingredients of Chinese medicines including luteolin, quercetin etc. could manage COVID-19 by targeting on AEC2 and 3CL protein and dampen inflammatory mediators without side effects and have achieved significant clinical efficacies (14). Our previous investigation found that luteolin can regulate the expression of inflammatory factors in macrophages and play an anti-inflammatory role (7), but whether luteolin can regulate the polarization of macrophages and its molecular mechanism is still veiled. In this research, bone marrow cells isolated from C57BL/6 mice were induced to differentiate into bone marrow-derived macrophages (BMDM) for investigating the effects of luteolin on the M1/2 polarization of BMDM and the expression of inflammatory factors so as to explore the underlying mechanism. Our present findings preliminary provided luteolin could be a future perspective for the natural anti-inflammatory agent in prevention and treatment of sepsis.

## MATERIALS AND METHODS

### Mice

6-week-old C57BL/6 mice (weighting 18-22 g) were provided from animal experimental center of affiliated hospital of integrated traditional Chinese and Western medicine, Nanjing University of Chinese medicine (Nanjing, China). Mice were maintained under specific pathogen-free conditions and in accordance with protocols approved by the National Institute of Health Guide for Care (Ethics Number: AEWC-20181019-53).

### Isolation and culture of BMDM

C57BL/6 mice were sacrificed by cervical dislocation, and dissected under immersion in 75% ethanol (V/V). The epiphysis was cut after tibia and femur were separated, and the bone marrow cavity was rinsed with sterile PBS until the bone became white. The bone marrow washing solution was filtered with a 70 μm mesh and transferred to a 50 mL centrifuge tube for cell collection. After lysis of red blood cells, bone marrow stem cells were resuspended in DMEM (Gibco, USA) containing 20 ng/mL M-CSF (R&D System, USA) on 10-cm petri dishes and renewed the medium every other day. After 7 days incubation, cells were labeled with F4/80-PE fluorescently conjugated antibodies (1.0 μL; eBioscience, USA) to identify the differentiation degree of mature BMDM by FCM (Millipore, USA).

### Cell viability assay

BMDM were plated in 96-well plates at a density of 2×10^4^ cells/well in 200 μL medium with confluence overnight. After that, the cells were exposed with LPS (20 ng/mL; Sigma, USA) plus IFN-γ (10 ng/mL; PeproTech, USA) or IL-4 (20 ng/mL; Pepro Tech, USA), and LPS plus IFN-γ-primed cells were administrated with luteolin (Sigma, USA) at 2.5 and 5.0 μmol/L for 24 h. Following treatment, cells were added with 20 μL MTT solution (5 mg/mL; Sigma, MO. USA). 4 h later, culture medium was removed and crystals were dissolved with 150 μL/well DMSO. The absorption values were measured at 570 nm.

### The morphology of polarized BMDM

BMDM in logarithmic stage were cultivated in 6-well plates, and incubated overnight followed by treatment with LPS (20 ng/mL) plus IFN-γ (10 ng/mL) or IL-4 (20 ng/mL). Simultaneously, the cells stimulated with LPS+IFN-γ were co-cultured with specified concentrations of luteolin for 24 h. The cell morphology was observed under an inverted microscope (Olympus, Japan). Cells and culture supernatants were collected for subsequent mRNA and protein analysis.

### Quantitative real-time PCR (qPCR)

BMDM were stimulated with LPS plus IFN-γ or IL-4 respectively, following dosing stimuli, cells were collected and total RNA was extracted by Trizol (Ambion Life Technology, USA) and reverse-transcribed into cDNA. With GAPDH as the internal reference, qPCR was carried out using SYBR Green Master Mix (Toyobo, Japan) according to primer sequences with Quant studio DX real-time quantitative PCR employed biosystems (Life Technologies, USA). Relative gene expression was calculated using the 2^-ΔΔct^ comparative method. Primer Sequences were obtained from Generay Biotech Co. Ltd. (Shanghai, China) and listed in Table 1.

**Table 1.**
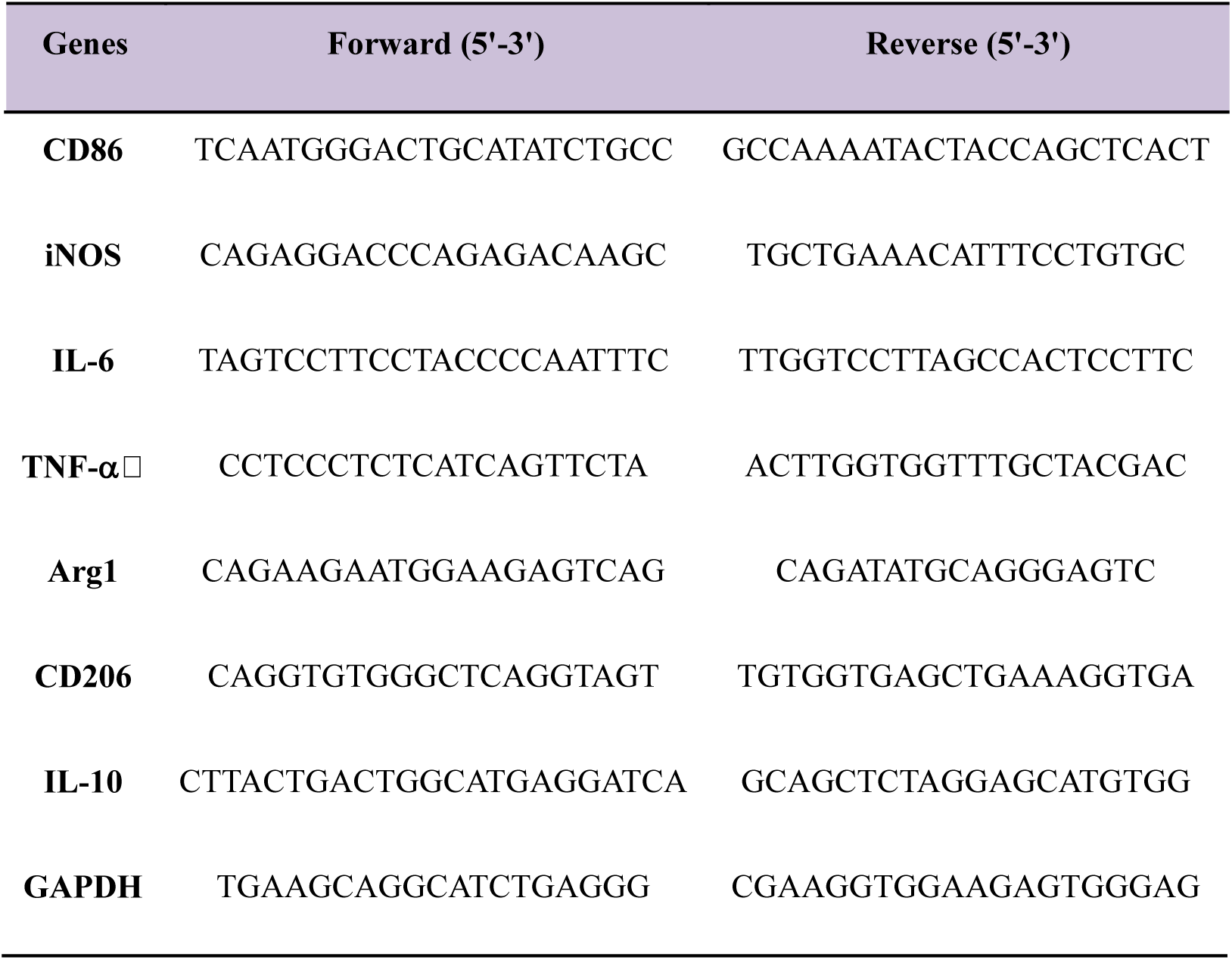
Primers for quantitative real-time PCR.

### ELISA for cytokines secretion

BMDM in exponential phase were inoculated in a six-well plate overnight. Successively, the drugs induced cell polarization and combined with different concentrations of luteolin for 24 h. The levels of cytokines in the supernatants were performed according to commercial ELISA kit (eBioscience, USA) instructions. Logistic fitting-curve for two of four parameters was used to calculate the concentration of cytokines.

### Flow Cytometric Staining of BMDM surface Markers

Totally, 5×10^5^ BMDM were resuspended in 100 μL PBS, then incubated with 0.5 μL anti-mouse CD16/32 blocking antibody (BioLegend, San Diego, USA) to avoid nonspecific binding in an ice bath for 20 min. Subsequently, cells were stained with anti-mouse FITC-CD11c (0.5 μL; BioLegend) or APC-CD206 (10 μL; BioLegend) and protected from exposure to light for 30 min at room temperature. After washing with PBS, the cells were fixed with 0.5 mL paraformaldehyde at 4°, and the mean fluorescence intensity (MFI) of membrane surface antigen CD11c or CD206 were analyzed by FCM.

### Protein extraction and immunoblotting

Collect cells and extract total protein for protein quantification by BCA. After 20 μg protein was subjected to SDS-PAGE and transferred to PVDF membrane (millipore, USA), the corresponding primary antibodies against p-STAT1-tyr^701^, p-STAT6-tyr^641^ (Cell Signaling Technology, USA) and β-actin (Sigma, USA) were applied at 4°C overnight. Then membranes were washed and incubated with HRP-conjugated secondary antibodies with shaken at room temperature for 30 min. Immunoreactive proteins were exposed and developed using ECL (Beyotime, China), and β-actin was used as an internal reference to calculate the relative expression of protein.

### Statistical analysis

The experimental data were presented as mean ± SD. One-way ANOVA followed by Tukey’s post-hoc test was used in the multiple comparisons. Analysis was performed using the GraphPad Prism 5.0 software (San Diego, CA, USA). *P*< 0.05 was considered statistically significant.

## RESULTS

### Mouse bone marrow cells differentiate into macrophages

The isolated mouse bone marrow cells were induced by M-CSF for 7 days, and FCM detected the specific marker F4/80 of mouse macrophages. The results showed that the purity of the differentiated macrophages reached 92.71%, indicating that the BMDM derived from mouse bone marrow were successfully cultured and could be used in subsequent experiments (Fig. 1).

**FIG 1.**
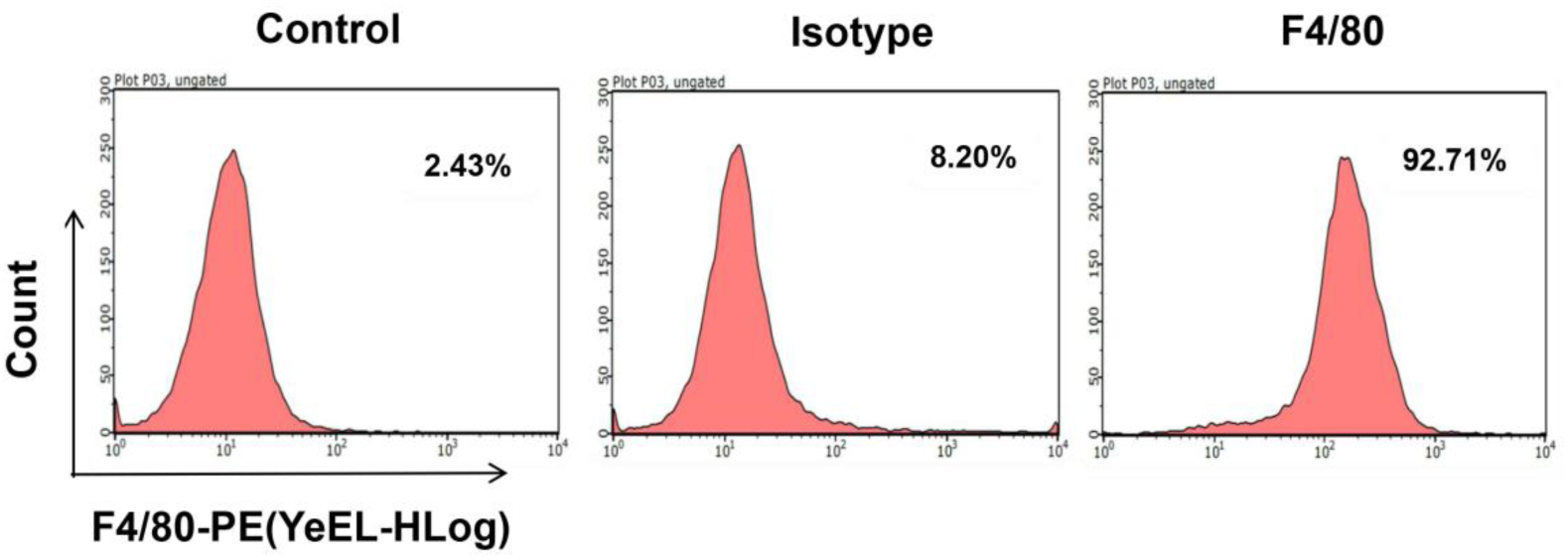
The differentiation proportion of BMDM. FCM detected the surface marker F4/80 of BMDM after induction for 7 days, and the positive rate was 92.71%.

### Effect of luteolin on BMDM viability

The non-cytotoxic doses of luteolin were evaluated via MTT assay to exclude contribution of anti-inflammatory potential of luteolin. Luteolin exhibited no significant impact on the LPS+IFN-γ-primed BMDM proliferation at concentration up to 5.0 μM at 24 h (Fig. 2 B). Therefore, non-cytotoxic concentration was chosen to assess the bioactivity of luteolin.

**FIG 2.**
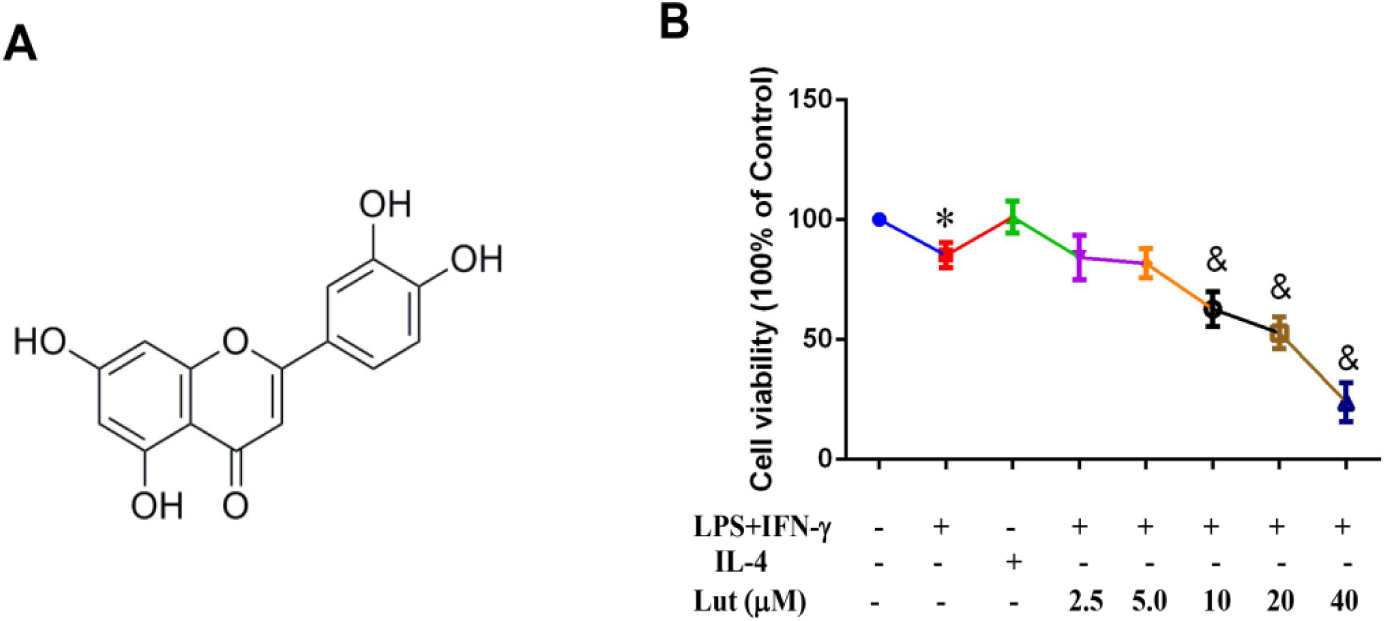
Effect of luteolin on LPS+IFN-γ-primed BMDM viability. A. Structure of luteolin. B. BMDM were primed with LPS+IFN-γ and contributed with indicated doses of luteolin for 24 h, then the cell viability was assessed by MTT assay. Data represented mean ± SD of three independent experiments performed in triple. Different symbols indicate a significant difference according to ANOVA and Tukey test. \**P* < 0.05 vs. control group (non-treated control); ^&^ *P* < 0.05 vs. LPS+IFN-γ-treated group.

### Morphology of polarized BMDM

Microscopically, BMDM showed typical morphology of macrophages, such as round, oval or irregular shape, with pseudopods and adherential growth (Fig.3. Control group). After being induced into M1-phenotype by LPS plus IFN-γ, the BMDM presented oval “Fried egg” appearance and pseudopodia extension (Fig.3. LPS+IFN-γ-treated group), while the M2-type BMDM induced by IL-4 were round and plump cytoplasm, accompanied by short pseudopodia (Fig. 3. IL-4-treated group). After various dose of luteolin contribution, M1 cells contracted slightly and pseudopodia became shorter (Fig. 3. LPS+IFN-γ-combined with 2.5/5.0 Lut-treated groups).

**FIG 3.**
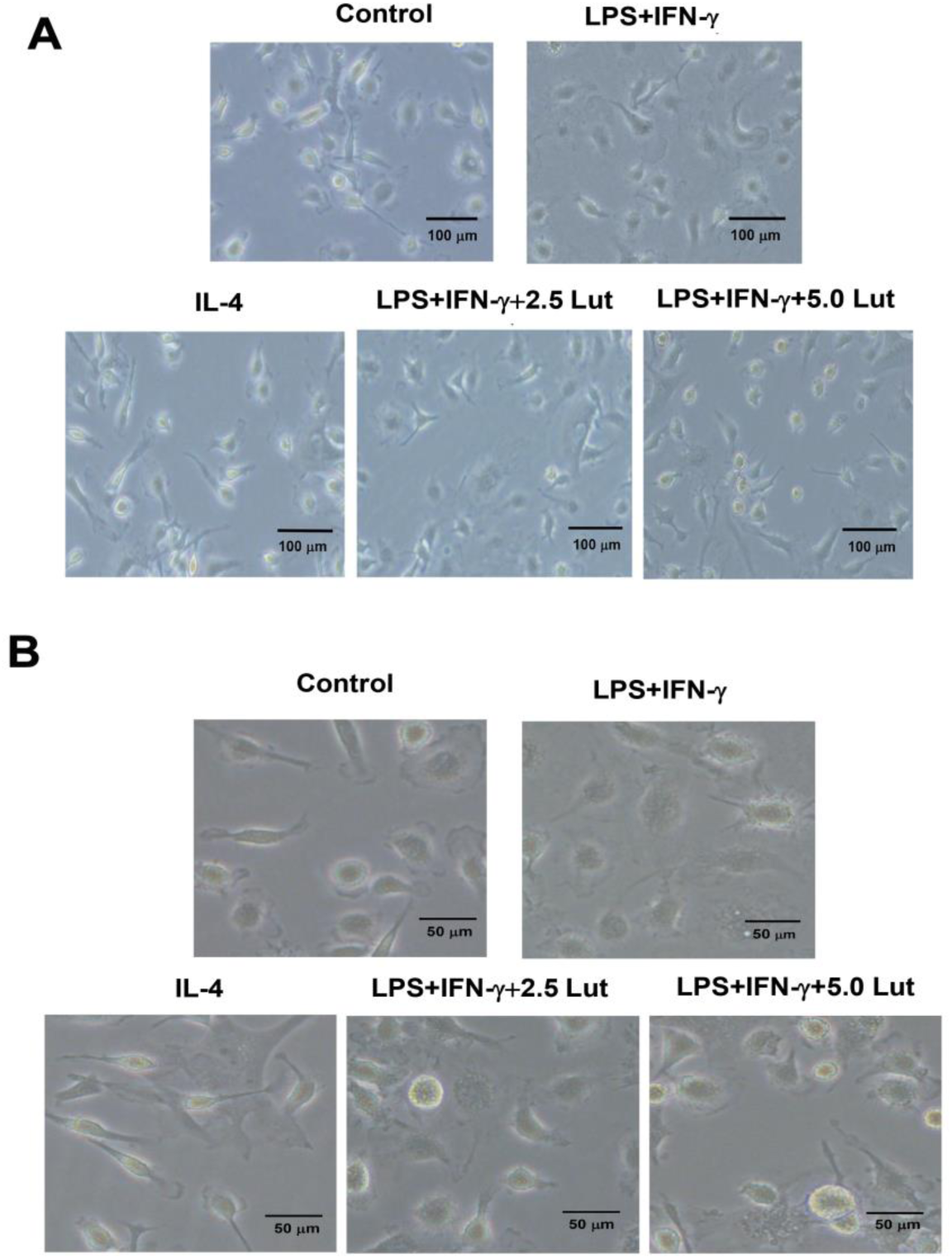
The morphology of polarized BMDM (A. 100×; B. 200×). BMDM were polarized with LPS+IFN-γ or IL-4, simultaneously, BMDM exposed to LPS+IFN-γ were administrated with luteolin for 24 h. Micrographs of BMDM were observed using bright field Olympus imaging system.

### Effect of luteolin on mRNA expression of inflammatory factors in polarized BMDM

The expression of M1-type and M2-type proinflammatory factors were detected by qPCR. Results implied that the expression of M1-type pro-inflammatory factors in M1-polarized BMDM such as CD86, iNOS, TNF-α and IL-6 was up-regulated; the expression of M2 type anti-inflammatory factors such as CD206, Arg1 and IL-10 in M2-polarized BMDM also up-modulated. This indicates that the polarization models of BMDM were successfully induced. When M1 cells were combined with luteolin, the expression of M1-type pro-inflammatory factors were decreased (Fig. 4), while the expression of M2-type anti-inflammatory factors were increased (Fig. 5).

**FIG 4.**
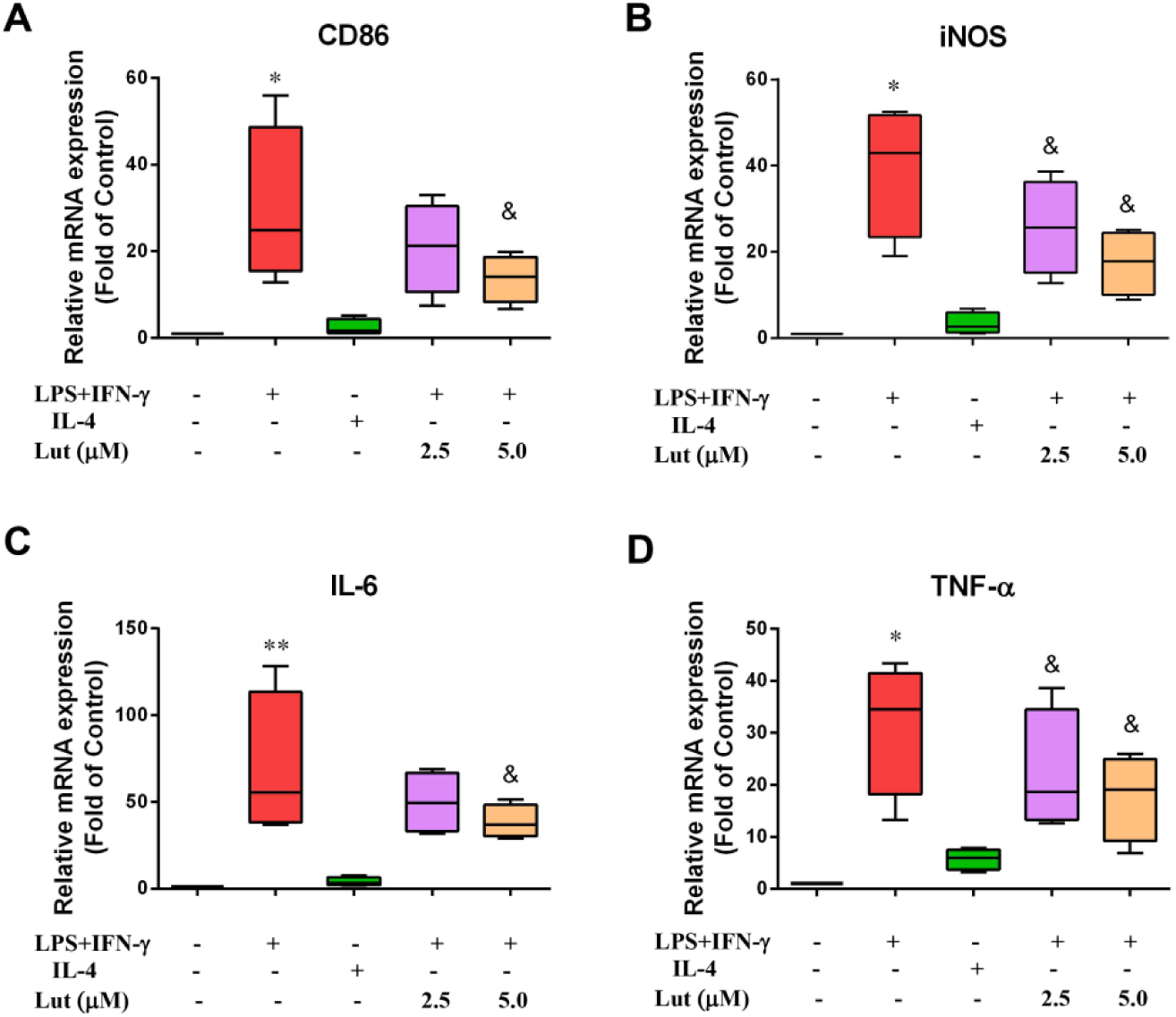
Effect of luteolin on the expression of M1-type pro-inflammatory factors in activated BMDM. The M1-type mRNA molecules were determined by qPCR with GAPDH as an internal control. BMDM were primed with LPS+IFN-γ or IL-4, and LPS+IFN-γ-treated-with indicated doses of luteolin for 24 h, the relative M1-type mRNA levels of CD86 (A), iNOS (B), IL-6 (C) and TNF-α -polarized macrophages reduced slowly. Data represented mean ± SD of four independent experiments performed in duplicate. Different symbols indicate a significant difference according to ANOVA and Tukey test. \**P* < 0.05, \*\**P* < 0.01 vs. control group (non-treated control); ^&^ *P* < 0.05 vs. LPS+IFN-γ-treated group.

**FIG 5.**
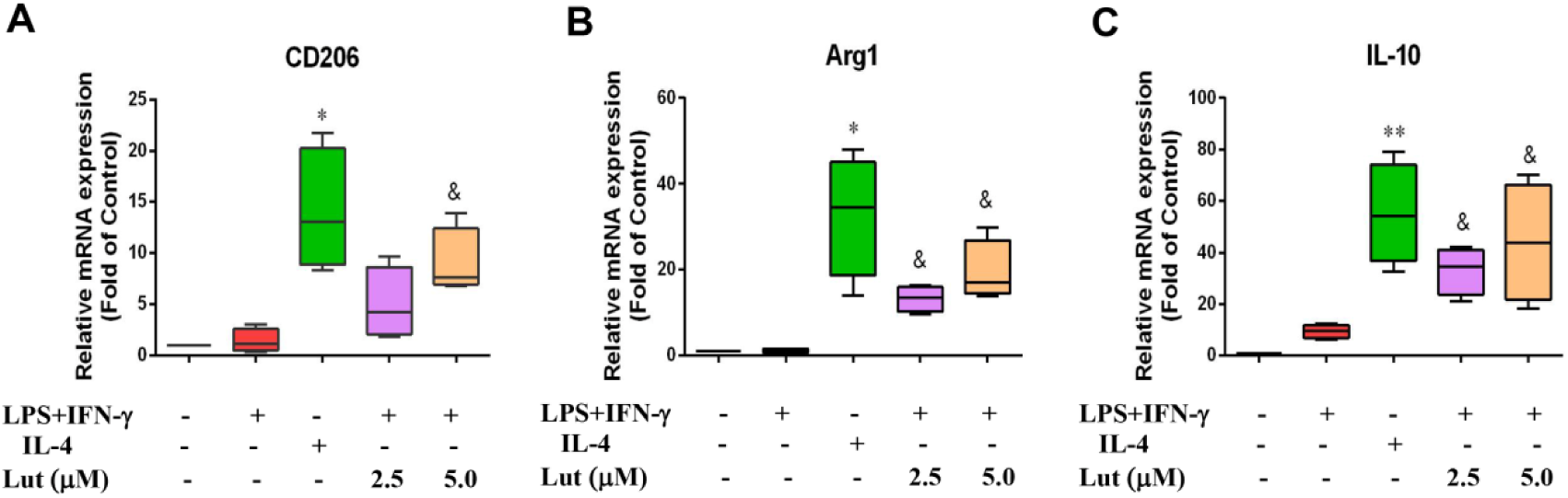
Effect of luteolin on the expression of M2-type anti-inflammatory factors in activated BMDM. The M2-type mRNA molecules were assessed by qPCR with GAPDH as an internal control. BMDM were primed with LPS+IFN-γ or IL-4, and LPS+IFN-γ-with indicated doses of luteolin for 24 h, the relative M2-type mRNA levels CD206 (A), Arg1 (B) and IL-10 (C) elevated gradually. Data represented mean ± SD of four independent experiments performed in duplicate. Different symbols indicate a significant difference according to ANOVA and Tukey test. \**P* < 0.05, \*\**P* < 0.01 vs. control group (without treatment); ^&^ *P* < 0.05 vs. LPS+IFN-γ-treated group.

### Effect of luteolin on inflammatory cytokine levels in polarized BMDM

To further explore the polarity skewing effect of luteolin in BMDM, IL-6 and IL-10 production were measured with ELISA. Pro-inflammatory cytokine IL-6 liberated by M1-polarized BMDM increased significantly, in the meantime, anti-inflammatory cytokine IL-10 secreted by M2-polarized BMDM also amplificated clearly, which was statistically different from the control group (without treatment). After various dose of luteolin challenged to M1-polarized BMDM, IL-6 released by M1-polarized BMDM lowered visibly, while IL-10 elevated obviously, compared with corresponding LPS+IFN-γ-treated group, there was a statistical difference (Fig. 6).

**FIG 6.**
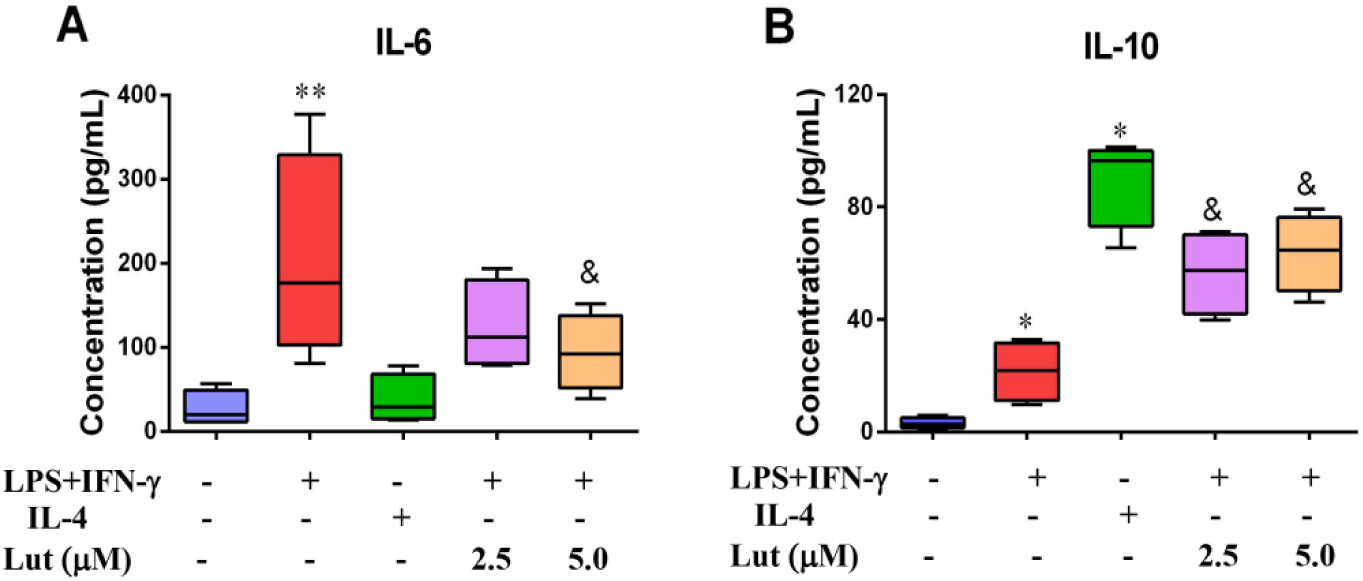
Effect of luteolin on IL-6 and IL-10 levels in polarized BMDM. BMDM were primed with LPS+IFN-γ or IL-4, followed by luteolin exposure for 24 h. Supernatants were harvested and levels of IL-6 (A) and IL-10 (B) secreted from the M-polarized BMDM were measured via ELISA. Data represented mean ± SD of four independent experiments performed in duplicate. Different symbols indicate a significant difference according to ANOVA and Tukey test. \**P*<0.05 vs. Control group (without treatment); ^&^ *P* < 0.05 vs. LPS+IFN-γ-treated group.

**FIG 7.**
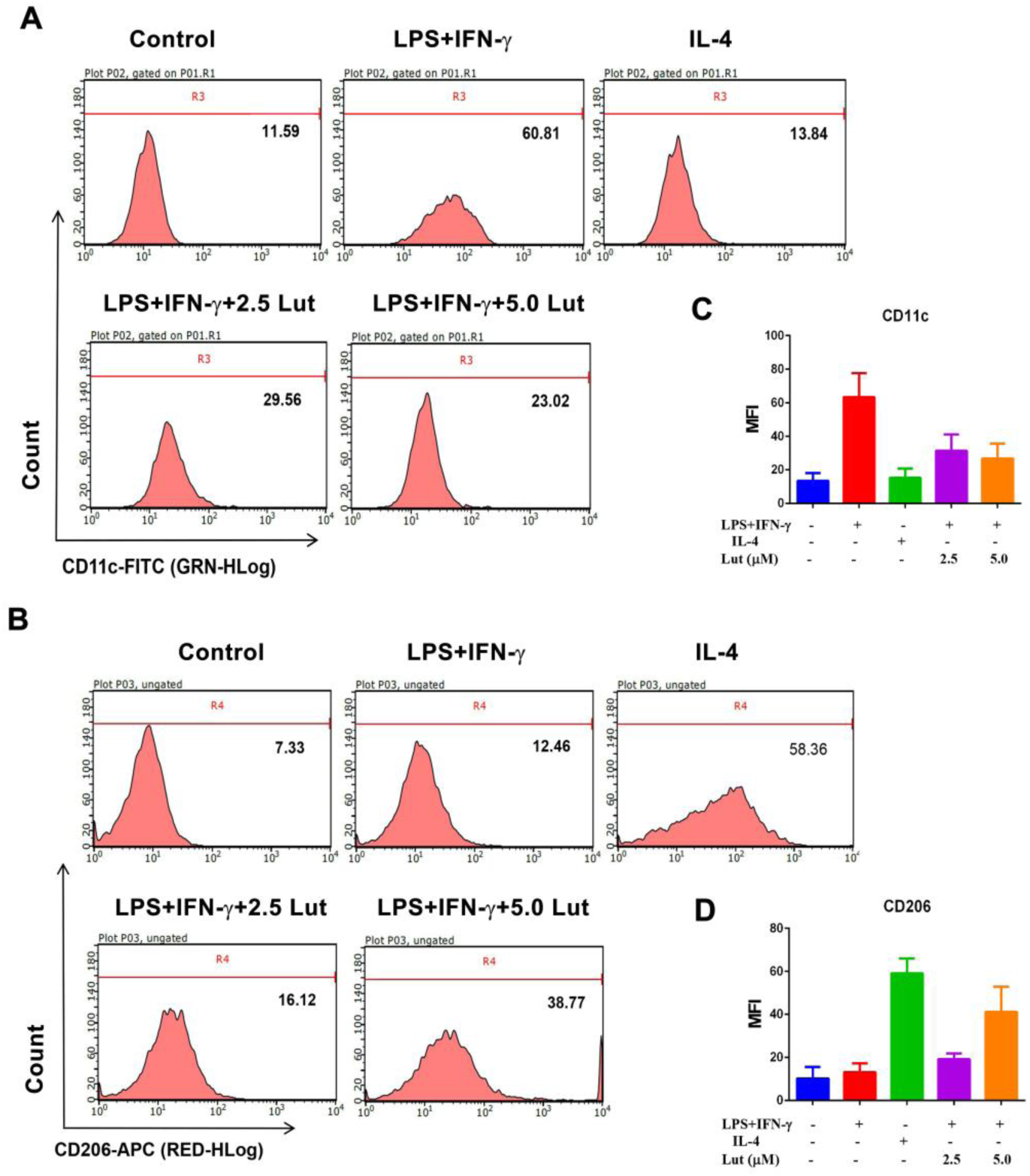
Effects of luteolin on the expression of BMDM surface markers CD11c and CD206. The BMDM were stimulated with LPS+IFN-γ or IL-4, and then luteolin treatment for 24 h, the expression levels of CD11c (A) and CD206 (B) protein on BMDM are presented as MFI as evaluated by FCM. The histogram presents the MFI of CD11c (C) and CD206 (D). Data represented mean ± SD of three independent experiments performed in triplicate. Different symbols indicate a significant difference according to ANOVA and Tukey test. \**P*<0.05 vs. Control group (without treatment); ^&^ *P* < 0.05 vs. LPS+IFN-γ-treated group.

### Effect of luteolin on the expression of surface markers on polarized BMDM

CD11c and CD206 are the surface marks of M1-polarized or M2-polarized BMDM, respectively. FCM results elevated that the MFI of CD11c (60.81) in M1-polarized BMDM was significantly enhanced compared with that of IL-4 treatment group (13.84) and Control group (11.59). The MFI of CD206 (58.36) in M2-polarized BMDM was also significantly amplified than that in LPS+IFN-γ alone treatment group (12.46) and Control group (7.33). Against this, luteolin treatment dramatically attenuated the CD11c MFI to 29.56 (LPS+IFN-γ+2.5Lut-treated group) and 23.02 (LPS+IFN-γ+5.0Lut-treated group) in M1-polarized BMDM, but gradually strengthened the CD206 MFI to 16.12 (LPS+IFN-γ+2.5 Lut-treated group) and 38.77 (LPS+IFN-γ+5.0 Lut-treated group) in M1-polarized BMDM in a concentration-dependent pattern (Fig.7).

### Effect of luteolin on protein pathway in polarized BMDM

STAT signaling proteins exert a vital role in macrophage polarization and the expression of inflammatory cytokines in sepsis. Immunoblotting assay and densitometry analysis of STAT proteins revealed that M1-polarized BMDM highly expresses p-STAT1 and lowly expresses p-STAT6, whereas M2-polarized BMDM lowly expresses p-STAT1 and highly expresses p-STAT6. Predominantly, after luteolin contribution, p-STAT1 expression was depressed while p-STAT6 was strengthened in protein pathway of M1-polarized BMDM (Fig. 8).

**FIG 8.**
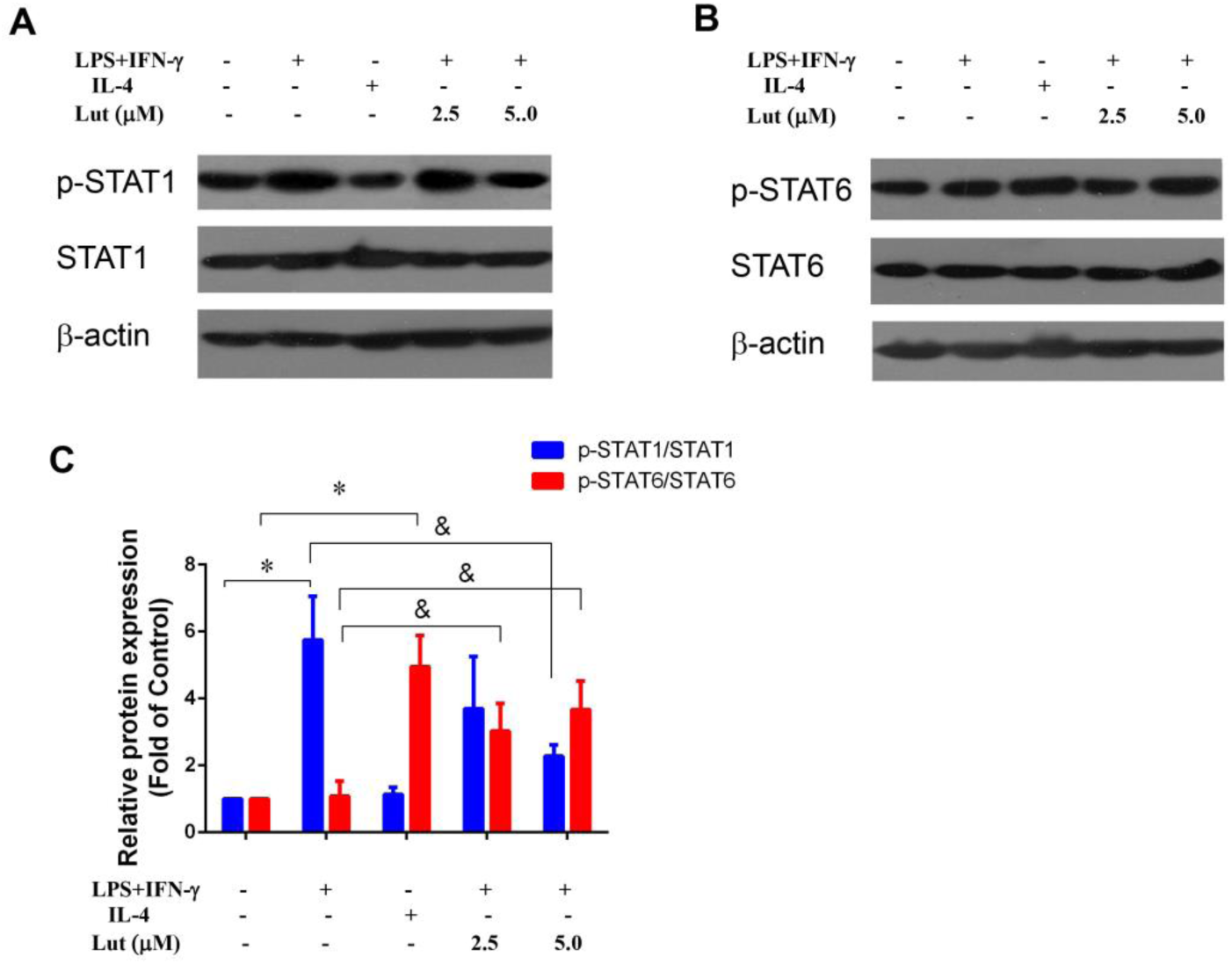
Effects of luteolin on the protein levels of p-STAT1/6 in polarized BMDM. The BMDM were primed with LPS+IFN-γ or IL-4, and then for addition of luteolin for 24 h. Total cell lysates were analyzed by immunoblotting for the indicated antibody, respectively. β-Actin was used as loading control. Representative immunoblots of p-STAT1 (A) and p-STAT6 (B); The relative protein levels of p-STAT1and p-STAT6 (D) by densitometric analysis. Data represented mean ± SD of three independent experiments. Different symbols indicate a significant difference according to ANOVA and Tukey test. \**P*<0.05 vs. Control group (without treatment); ^&^ *P* < 0.05 vs. LPS+IFN-γ-treated group.

## DISCUSSION

Infectious diseases are the leading cause of death, and the infection severity is due to an exaggerated activation of macrophages and cytokine storm (15). LPS, the main component of endotoxin, is the outer membrane structure of cell wall of Gram-negative bacteria which can bind to toll-like receptor 4 (TLR4) on macrophage surface to induce M1 polarization and secrete pleiotropic cytokines. IFN-γ can synergy with LPS to further activate cells, secrete excessive cytokines, cause SIRS, and severe cases cause sepsis and MOF (7). Therefore, it is particularly important to regulate macrophage polarization, avoid excessive activation of M1 macrophages, reduce inflammation and promote tissue repair. BMDM are suitable cell models for studying macrophage polarization. In this investigation, bone marrow cells derived from femur of C57BL/6 mice were stimulated to develop and differentiate into mature BMDM by M-CSF. LPS and IFN-γ stimulated BMDM to undergo M1 polarization, and M1-type pro-inflammatory factors including iNOS, TNF, IL-6 and surface markers CD86 and CD11C were up-regulated; IL-4 stimulated BMDM M2 polarization, and M2-type anti-inflammatory factors including Arg1, IL-10 and CD206 were up-regulated, indicating successful induction of M1/2 polarization in BMDM. After luteolin contribution, the M1-type pro-inflammatory factors decreased and the M2-type anti-inflammatory factors increased evidently in M1-polarized BMDM. Concurrently, the protein pathway p-STAT1 expression was down-regulated and p-STAT6 expression was up-regulated. It suggests that luteolin may modulate the phenotype polarization of BMDM through the inhibition of p-STAT1 and the activation of p-STTA6, transforming it from pro-inflammatory M1-type to anti-inflammatory M2-type, thereby reducing the expression of inflammatory mediators and alleviating inflammation to maintain the stability of the microenvironment.

Cell polarization is regulated by various signaling molecules or transcription factors. STATs are one of the pivotal signal transduction pathways and widely involved in the process of cell activation, apoptosis, inflammation and immune regulation (16). Studies by Sodhi et al. (17) and Zhou et al. (18) showed that IL-6 and IFN-γ released by LPS-polarized M1 macrophages can promote the expression of STAT1 protein, and that IFN-γ can also motivate STAT1 by binding to its receptor, and simultaneously, the activated STAT1 can further provoke the levels of TNF-α, IL-1β and iNOS in macrophages. iNOS, is a signature of M1-polarized macrophages, which responsible for nitric oxide (NO) production when cells are stimulated by IFN-γ or LPS, and excessive NO causes oxidative stress and inflammatory damage (19). Both CD86 and CD11c are surface markers of M1 macrophages. CD86 is a B7 costimulatory molecule that stimulates the activation of antigen-presenting cells to secrete more pro-inflammatory factors. In the meantime, the level of CD86 can reflect and positively correlate with the level of cytokines such as IFN-γ and IL-12, while IL-10 can hinder the level of CD86 (20-210). CD11c is often coupled with CD18 and binds to bacterial LPS, which activates CD4^+^T cells to proliferate and differentiate into Th1 cells and secrete massive TNF-α, IL-6 and IL-12 to trigger inflammatory cascades (22). Two other crucial cytokines IL-6 and TNF-α, generated abundantly by IL-1β stimulation or autocrine from activated “mononuclear-macrophage system”, are elevated not only in bacterial infection but also during viral infection (23-24). More importantly, they are most strong pro-inflammatory agent causing “cytokine storm”. Studies have shown that patients with severe COVID-19 characterized by a “cytokine storm” inexorably exhibited high levels of IL-6 and TNF-α in serum, and IL-6 or TNF-α antagonist seems to be very promising for severe COVID-19 cases (25-26).

To our knowledge, IL-4 or IL-13 can induce M2-type polarization of macrophages, and M2-type anti-inflammatory factors such as IL-10, Arg1 and CD206 are up-modulated. In this regard, IL-4 binds to its receptor to activate JAK to further phosphorylate STAT6 and enhance the Arg1 activity. Arg1 and iNOS are important hallmarkers of M2/M1 type macrophage polarization, respectively. Under normal circumstances, the activities of Arg1 and iNOS are strictly regulated by macrophages and maintain a dynamic equilibrium. When M2 polarization occurs, Arg1 competes for iNOS to decompose substrate arginine, thus benefit for tissue regeneration. Moreover, Arg1 is also inseparable from M2 macrophage properties in playing immune memory function to eliminate infectious agents. CD206, so called mannose receptor, is a membrane surface marker of M2 cells, which can specifically recognize antigens to clear pathogens, promote angiogenesis and repress immune response (27). Another M2-type anti-inflammatory factor, IL-10, on the one hand, enhances the sensitivity of macrophages to IL-4 and IL-13 by increasing the abundance of IL-4 receptors on the macrophage surface, which contributes to M2-type polarization of macrophages. On the other hand, it can synergize with IL-4 to inhibit pro-inflammatory cytokines IL-1β and TNF-α to reduce inflammation. In the light of preliminary data, IL-10 displayed higher levels in patients with sepsis and serious COVID-19 (28). All these indicate that when inflammation is motivated, an intricate network is formed between the pro-inflammatory mediators and activated STAT1, eliciting “inflammatory storm.” Nevertheless, upon luteolin contribution, a complex network is also formed between anti-inflammatory mediators and activated STAT6, further facilitating the expression of anti-inflammatory factors which resist the formation of pro-inflammatory factors and alleviate inflammation accordingly. Herbal compound Physalin D can repolarize M1 toward M2 polarization in BMDM through STAT1 suppression and STAT6 activation (29), which consist with our study.

Altogether, BMDM polarization mechanism is complex and involves many protein pathways. Only by actively exploring the regulation of BMDM polarization and maintaining the balance of inflammatory mediators can maintain the physical stable. In this investigation, LPS/IFN-γ induced M1 polarization and IL-4 induced M2 polarization of BMDM. After being treated with herbal compound luteolin, the M1 polarized BMDM showed lowered M1-type pro-inflammatory factors and elevated M2-type anti-inflammatory factor, and that signaling protein p-STAT1 was down-regulated and P-STAT6 was up-regulated. That is, the macrophage population underwent a transformation from a pro-inflammatory M1-phenotype to an anti-inflammator M2-phenotype. Simultaneously, inflammatory factors analogously altered from pro-inflammatory to anti-inflammatory. In light of these findings, our research provide a novel insight into the role of luteolin to be a candidate for controlling macrophage phenotype to treat infectious disease.

## ACKNOWLEDGEMENTS

We acknowledge the support of their staff and Facility of department of Endocrine and Metabolism in accomplishing this work. We also thank Dr. C. M. (Cao Meng) and Z. J. (Zhou Jing) for carrying out exploratory experiments and valuable advice. W. S. (Wang Shuxia) wrote the manuscript. Moreover, L. X. (Li Xingjia), M. X. (Mao Xiaodong), Y.W. (Yang Wanwei) and C. G. (Chen Guofang) edited this manuscript and analyzed data.

This work was supported by National Natural Science Foundation of China (grant number 81673945; 81471010) and the Project of Jiangsu Branch of China Academy of Chinese Medical Sciences (FY201809).

The authors declare there are no competing interests.

